# A Novel Restraint Device to Improve Safety and Efficacy of Blood Collection During Non-Terminal Sampling of Bats

**DOI:** 10.1101/2022.03.22.483499

**Authors:** Clint N. Morgan, Matthew R. Mauldin, Jeremy Jones, Brian Collier, Yoshinori Nakazawa

## Abstract

There are a variety of blood collection techniques described in the literature for unanesthetized bats, which typically require multiple sharps (e.g., needles, lancets, etc.), competent animal handling for prolonged periods, and usually involve two individuals. With the challenges inherent to non-terminal sampling of blood from bats, as well as the growing need for the use of this technique across multiple disciplines and industries, an improved blood collection method is needed. We report the creation of a bat restraint device specifically designed for a single individual to safely collect blood from anesthetized or non-anesthetized bats. The utility of this restraint device is multifaceted, serving as a safety measure for both animal and handler, as well as increasing the efficiency of blood collection. The restraint device was tested during two laboratory bat studies, Afterwards, the users of the restraint device were provided with a 10-question survey questionnaire to record their opinions on its usage. In total 80% of responses were considered positive, 15% considered neutral, and 5% considered negative. Survey questions that all participants responded to positively when in comparison to the traditional method of blood collection from bats include “easier to perform”, “safer to bats”, and “safer to the individual”. While using the restraint devices during the laboratory studies, no needle sticks, bites, or scratches to laboratorians occurred, and no observable health issues or complication due to blood collection in the bats bled using the restraint devices.

## Introduction

Bats are the second largest order of mammals, and the research need for disease surveillance in bats continues to grow with the influx of both emerging infectious diseases affecting bats, and bat-associated high-consequence zoonotic diseases such as lyssaviruses, filoviruses and coronaviruses [1,2]. Based on the current whole-genome sequence analysis data, Severe Acute Respiratory Syndrome Coronavirus 2 (SARS-CoV-2) that caused a global pandemic beginning in 2019, is most similar to a SARS-CoV-like coronavirus (RaTG13) isolated from bats [3,4]. In response, there has been increased national and international interest in conducting monitoring and surveillance of susceptible animals for SARS-CoV-2, and strengthening the ability for early detection of emerging and zoonotic diseases in bats and other animals [5]. There has also been an increase in bat conservation efforts across the world, and disease surveillance for non-zoonotic bat diseases such as White Nose Syndrome, caused by an infection of the fungal pathogen *Pseudogymnoascus destructans* [6]. This fungal pathogen has resulted in the collapse of North American bat populations since its introduction from Eurasia as early as 2006 [7]. Serological surveillance is commonly used by wildlife disease researchers to assess the presence of antibodies to a specific pathogen in the blood of wildlife populations. Serosurveillance gives researchers insight into the prevalence and distribution of a pathogen in a population, and has been used with success to detect pathogens in regions previously undetected by other means [8,9]. Effective, safe, and efficient non-terminal blood collection is essential for successful serosurveillance, and we herein describe an improved method for non-terminal blood collection from bats specifically, using a novel restraint device.

Bats consist of two major taxonomic groups (Suborder), Microchiroptera (hereafter termed Microbats), which occur worldwide and are the most diverse, and Megachiroptera (hereafter termed Megabats), which are commonly referred to as “fruit bats” are restricted to the Old World. Blood collection from Microbats is typically more challenging than from Megabats as their veins are smaller and can be difficult to puncture with a small gauge needle. A variety of non-terminal blood collection techniques for bats are described in the literature, with all typically following the recommendations by the American Society of Mammalogists in field settings [10], or guidelines established by the National Research Council in laboratory settings [11]. Blood collection from bats typically entails using a 25-30 gauge needle (depending on the size of the bat) to puncture a peripheral vein, typically utilizing the cephalic vein (on the antebrachial wing membrane) or saphenous (interfemoral) vein [12,13]. Following venipuncture, the blood is allowed to pool, and is collected via glass or plastic capillary tube. This technique typically requires multiple sharps and relies on the competence of researchers to manually restrain the bat, and to don appropriate personal protective equipment in efforts to avoid bites and scratches.

Given the inherent difficulty of this technique, it is typically required that at least two individuals perform the blood collection, with one individual manually restraining the bat with leather gloved hands, and the other individual without leather gloves (for increased dexterity) handling the needle and collecting the blood. This “traditional” method requires competent animal handling for prolonged periods. If a researcher has many bats to sample, this traditional method can lead to mental and physical fatigue which increases the risks of adverse events occurring.

Bat blood collection can be performed while bats are either anesthetized or unanesthetized, with anesthesia being the method preferred by some researchers to avoid the potential for bites and scratches during prolonged restraint. Use of anesthesia may be used in efforts to limit the stress on the animal or to increase safety to the handler; however, it also requires expensive equipment and can lead to health complications if anesthetic gas is not delivered (or recovered) competently. Blood collection from unanesthetized bats is typically a preferred field method as it requires less specialized and expensive equipment. However, blood collection from unanesthetized bats typically comes with the increased risk to the researcher of bite or scratch, increased duration of blood collection which can be physically taxing to both animal and researcher, and limits the overall quantity of bats that can be sampled due to time constraints. In efforts to alleviate the stress on both animal and researcher, as well as decrease the duration and level of difficulty in blood collection, an evaluation of the bat blood collection method in the literature was conducted in effort to develop an alternative method for blood collection in bats, e.g., the use of restraint devices.

Restraint devices are commonly used for blood collection from rodent species and a variety of other taxa in laboratory settings. The typical restraint device is intended to safely house the animal, limit its movement, and provide access to the target area for blood collection. Another large benefit of using restraint devices is that the work can typically be performed by one individual which can increase productivity, efficiency, and safety, especially in high-containment laboratory settings. Improvised restraint devices have been described for Microbats [13], however there are no such devices documented to achieve safe blood collection from unanesthetized bats, or Megabats. A restraint device for blood collection of bats, if safe and effective, could prevent bites or injuries to the bat and researchers, limit the movement of the bat, allow easy access to the common peripheral veins used for blood collection (saphenous and cephalic veins), and decrease the number of individuals needed for the blood collection activity. Given that there are situations where anesthesia of the bat is preferred, a restraint device that doubles as an induction capsule would be preferred.

In an effort to meet these safety and efficiency needs, a restraint device for the safe blood collection of bats was developed and tested during two laboratory bat study with two species of Microbat, Big Brown Bat (*Eptesicus fuscus*) and Brazilian Free-tailed bat (*Tadarida brasiliensis*). Two models of restraint device were developed to best fit the size differences each species, in efforts to demonstrate that this restraint platform can be expanded and developed for any bat species. Following this pilot study, the researchers and technicians who had used the restraint device were provided with a survey questionnaire to record their user experience and attitudes toward the restraints when compared to the traditional method of blood collection in bats.

## Materials and Methods

### Bat Restraint Device Description & Use

The bat restraint device, as shown in Figure 1, comprises a base designed to sit atop a flat work surface, with neodymium magnets to allow for affixing to a metal surface (Fig 1a.), a hollow sliding plunger designed to secure the bat into position within the capsule without limiting airflow, additionally a ¼” ID vinyl hose can be secured on the end to be adapted for use with an anesthetic vaporizer (Fig 1b.) a sliding snap-ring that affixes to the capsule allows the bat forearm/wing to be pulled out of the capsule slot and secured in place for safe wing vein collection (Fig 1c.). The top portion of the bat capsule is constructed of a clear vinyl allowing for visibility within the capsule (Fig 1d.). The bottom portion of the bat capsule secures firmly to the sliding rail on the base and is constructed to limit movement in all directions (Fig 1e.). The two halves of the capsule can be separated quickly using the quick-release tabs to release the bat (Fig 1f.). The device is a 3D-printed non-porous nylon material, designed for durability and to be easily dismantled for cleaning purposes. The capsule within which the bat is placed is elevated, allowing space and access underneath the bat to aid in the blood collection. The dimensions of this restraint device are easily scalable to accommodate the extreme range of size differences among bats (e.g., Microbats vs Megabats).

**Figure 1.**
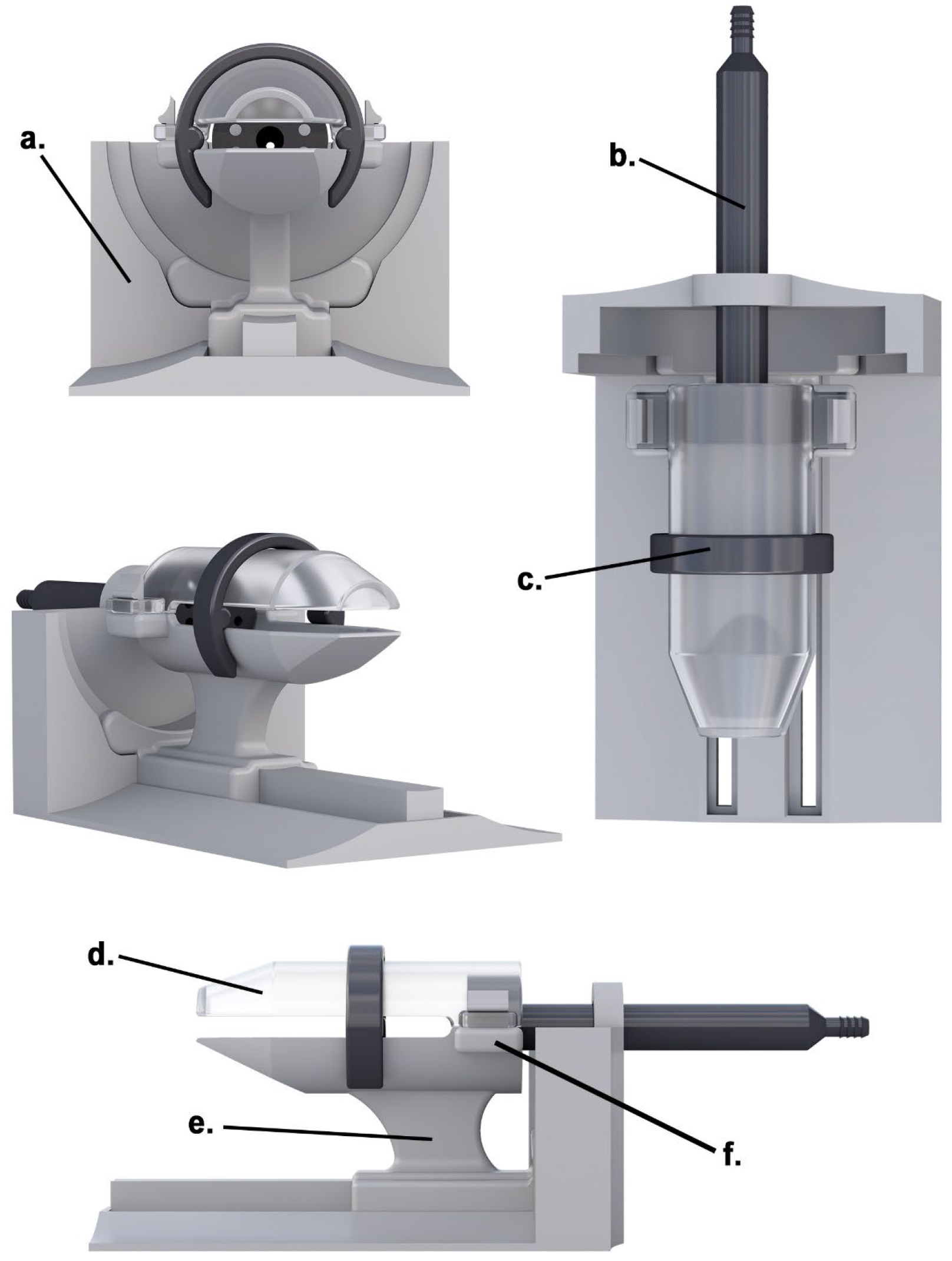
Example of the bat restraint device to safely restrain bats during blood collection; a. Restraint base provides stability and attachment rail/slot for capsule, b. hollow sliding plunger used to secure bat into capsule, c. sliding snap-ring allows bat wing to be secured out of the capsule slot for wing vein collection, d. Top portion of capsule (clear vinyl), e. Bottom portion of capsule with integrated rails on base for stability in all directions, f. Quick-release tabs, allowing capsule to be opened/closed as necessary.

A bat’s natural escape behavior is to crawl (or fly) upwards, also they display backwards “searching” behavior with their hind feet searching for somewhere to grasp [14]. This natural behavior is the preferred method to “load” the bat into the restraint device. This procedure is described as follows. While handling the bat with leather-gloved hands, place the caudal end (legs and uropatagium) into the capsule, then tilt upwards allowing the bat to move backwards into the capsule ensuring the wings are folded in along the body. The plunger can be subsequently inserted. Alternatively, the bat can be gently placed on the bottom half of the capsule facing the plunger, and the top half placed above and pressed gently to lock the quick-release tabs into place. The sliding snap-ring can then be slid into the capsule slot. To collect from the cephalic wing vein, the researcher can remove the snap-ring, grasp and stretch the wing through the slot, then slide the snap-ring upwards and into place under the shoulder of the bat which will safely preclude the humerus (and wing) from re-entering the capsule (Fig 2A). To collect blood from the saphenous vein, the researcher can gently grasp the hindfeet/uropatagium and pull through the slit at the end of the capsule (Fig 2B).

**Figure 2.**
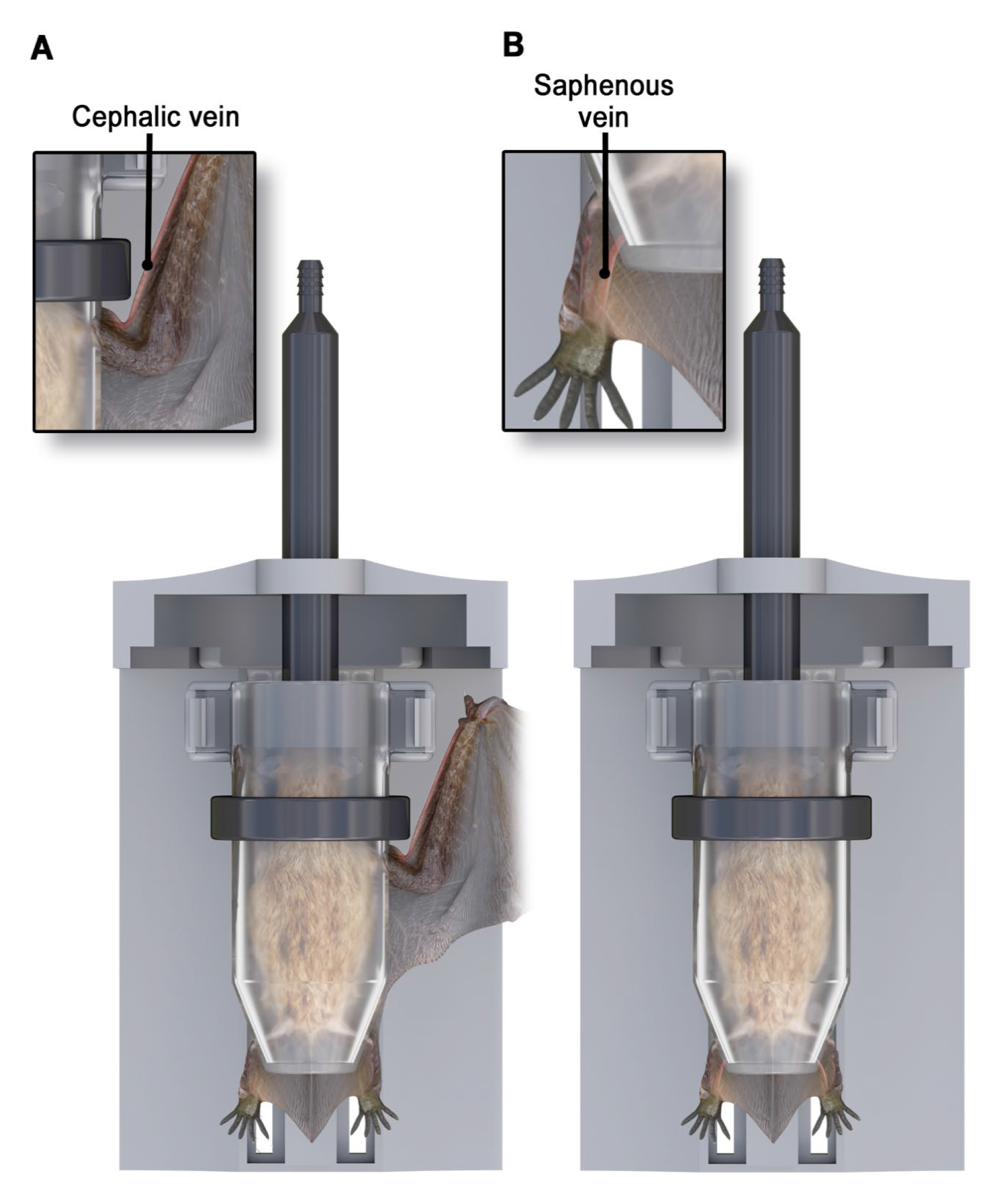
Figures display the position of the bat in the restraint devise for blood collection from the cephalic vein of the wing and the saphenous vein of the uropatagium. A. the wing and forearm can be drawn out of the capsule to access the cephalic vein, located on the leading edge of propatagium; note the position of the sliding snap-ring to lock the wing out of the capsule. B. the feet, tail and uropatagium can be drawn out of the end of the capsule to access the saphenous vein.

### Blood Collection

All laboratory personnel had prior experience with handling bats and peripheral vein blood collection, and were trained appropriately to use the restraint devices prior to blood collection. Each bat was manually restrained and the surface of the skin above the selected peripheral vein was disinfected with an alcohol-prep pad and allowed to air dry. Venipuncture was performed with a small gauge needle (27-25 gauge) and blood was allowed to pool into a droplet on the skin [15]. A 70 µL heparinized glass capillary tube was used to contact the blood droplet and through capillary action the blood is drawn up into the tube. Once the target blood volume was acquired, a styptic powder (Kwik Stop Styptic Powder, MiracleCorp Products, Dayton, OH, USA) is applied with pressure (pinch) to the puncture site until bleeding ceases. The bat was then released from the restraint and provided with fluids (water or fruit juice) orally to re-hydrate.

### Laboratory Testing

Two common North American bat species were used to test the safety and efficacy of the bat restraint device, the Brazilian free-tailed bat (*T. brasiliensis*), and the Big Brown Bat (*E. fuscus*). Testing of the restraint devices was opportunistically conducted over the course of June 2019 – Feb 2021 and approved by the Center for Disease Control and Prevention’s (CDC’s) Institutional Animal Care and Use Committee (IACUC) protocol numbers 3019NAKBATL and 3128HUTBATL. These bats were co-housed in a laboratory setting, and blood was collected at intervals determined by the respective study objectives, and not solely for the testing of restraint devices. Bats were weighed prior to each blood collection and a blood sample no larger than 1% of the body weight was collected [11].

### Survey Questionnaire

A ten-question survey questionnaire was developed and distributed to those that fit the inclusion criteria. The questionnaire solicited responses intended to compare multiple attributes of the traditional method and the restraint device method of blood collection (Table 1). Inclusion criteria for the survey participation included only individuals who had bat blood collection experience with both the traditional method (either in the lab or in the field), and also had used the restraint device on at least 2 occasions during the laboratory testing. Each participant’s survey answers were kept anonymous. Responses were compiled and analyzed for agreement among participants.

**Table 1.**
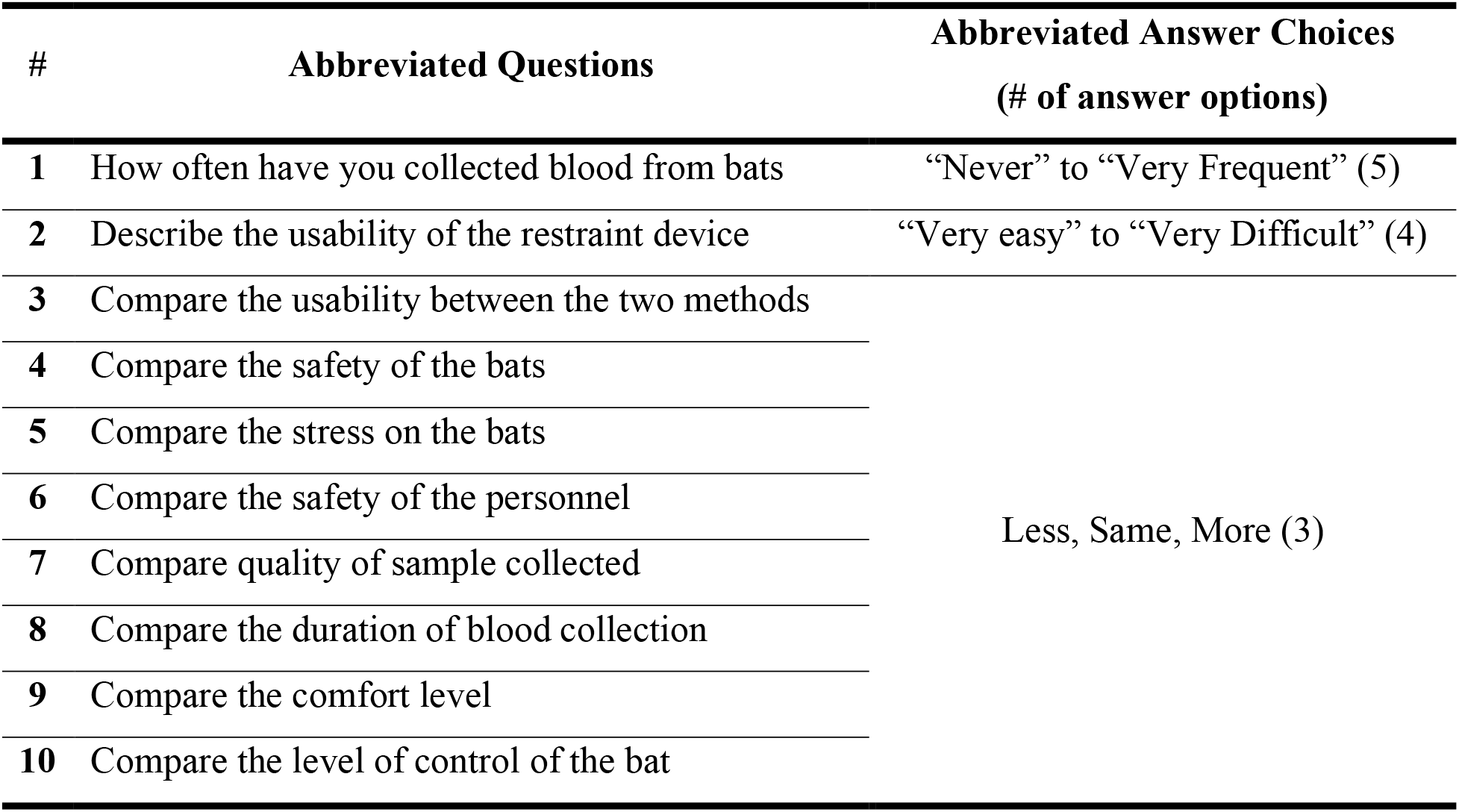
Abbreviated survey questionnaire questions. Comparisons solicited are between the use of the restraint devices for blood collection, versus the traditional method (manual restraint).

## Results

During the laboratory testing of the restraint devices, researchers and technicians collected blood on 9 occasions, and a combined total of 54 times from *T. brasiliensis* and 56 times from *E. fuscus* (including resampling of individuals). Throughout the laboratory studies the restraint devices were developed and improved upon, the final optimized design (dimensions scalable for different sized bats) is shown in Figure 1. While using the restraint devices during the laboratory studies, no needle sticks, bites, or scratches to laboratorians occurred. Additionally, there were no observable health issues or complication due to blood collection in the bats bled using the restraint devices.

In total, 6 participants were solicited and asked to complete and return the survey anonymously. Five participants completed and returned the survey questionnaire, with credentials ranging from veterinarians, researchers, and animal technicians. In the survey the participants established that they had either “occasionally” (n=3) or “somewhat frequently” (n=2) collected blood from bats. Without comparing the restraint devices to the traditional manual restraint method, the participants stated that the restraint device was “somewhat easy to use” (n=3) or “very easy to use” (n=2). In total 80% of survey responses (Fig. 3) were considered positively in favor of the restraint device (e.g., safer for the bat), 15% considered neutral (e.g., as stressful to bats), and 5% considered negative (e.g., longer duration of sample collection). The majority of questions related to the use of the restraint device when compared to the traditional method were considered positive, in favor of using restraint devices. Three of the comparison questions had 100% agreement among responses (ease of use, safety to bat, safety to researcher), 2 questions had 80% of participants positively favoring the restraint device (comfort level, control of bat), and 3 with 60% of participants positively favoring the restraint device (Stress level of bat, quality of sample, duration of collection). The questions that had the lowest agreement among participants include amount of stress likely experienced by the bats, the quality of blood sample collected, and the overall duration of blood collection. Only one question referring to the duration of time during the blood collection had participants negatively favoring the restraint device, with 2 participants stating that it takes “more time” when using the restraints.

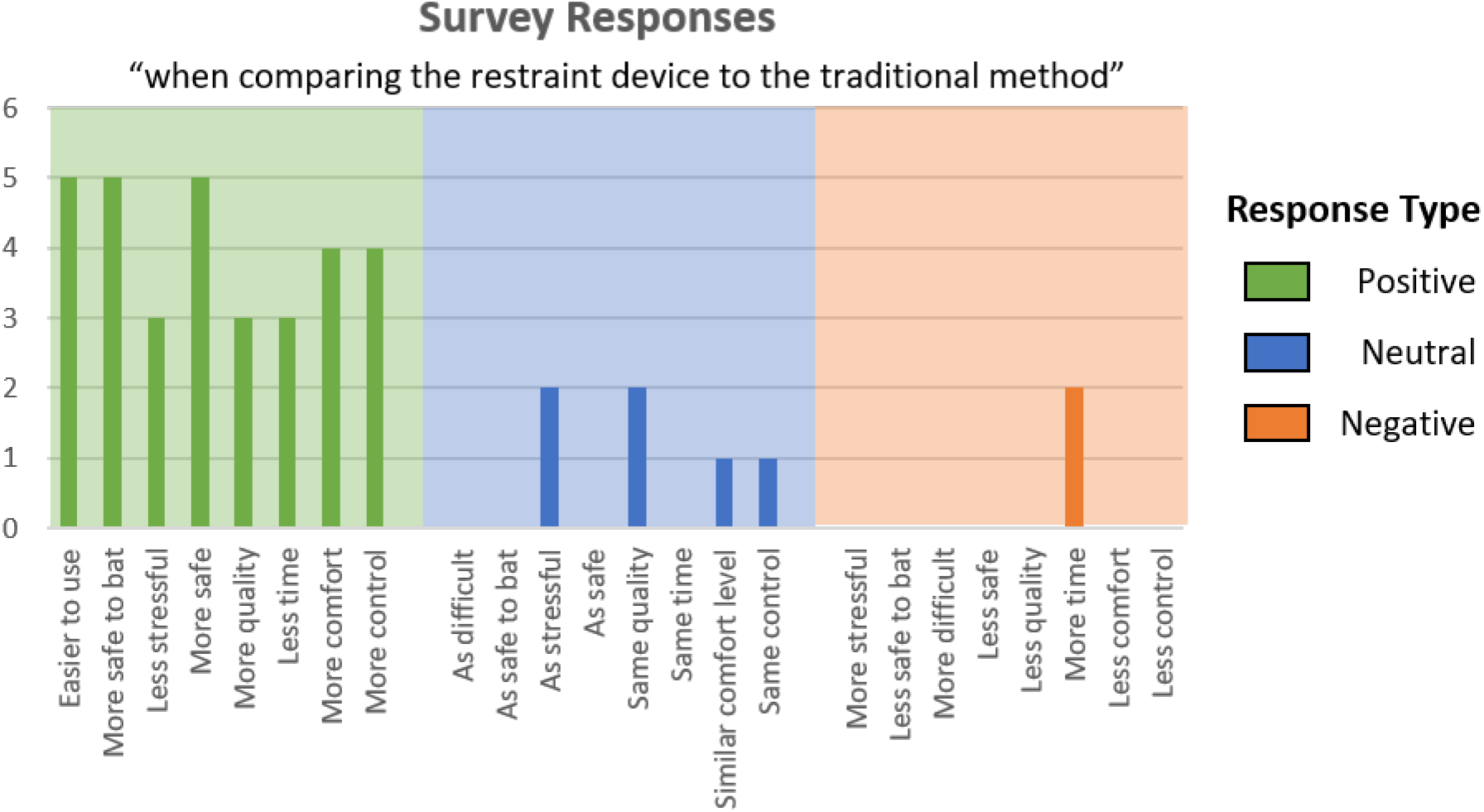

## Discussion

Serological studies of bats are expected to become more common in the future due to the involvement of these animals in the circulation of emerging pathogens of interest to public health and bat conservation. The use of restraint devices to increase the safety and efficiency of non-terminal blood collection will be an improvement over the current technique of manual restraint. The survey results of the restraint users are clear in that there is multifaceted benefit when using a restraint device. However, this was measured with a convenience sample of participants, and will need more extensive use and evaluation to fully assess the usage of a bat restraint device. The main perceived benefits to using the restraints (100% agreement among participants) were increased safety to the bat and the individual, as well as being overall less difficult. Increasing safety to bats and researchers alone are large benefits, as bats when restrained by hand can wriggle excessively, delivering defensive bites, or scratch an unprotected handler [15]. Although certain sample collection strategies and bat species may not call for the use of these restraints; for example, venipuncture of large peripheral veins of some Megabats may be more efficient without the use of a restraint device. However, in this same scenario the restraint devices may provide the benefit of allowing for just one individual to collect blood, which may increase the productivity of the sampling effort.

This restraint device is not intended for use in terminal sampling and should be used only for nonterminal blood collection. Terminal blood collection, blood collection under anesthesia immediately followed by euthanasia, is commonly conducted in laboratory environments following experimental infection with a pathogen, or at the termination of a study. Terminal blood collection is also performed in the field, however permitting agencies often preclude these activities due to conservation or population management concerns, or when working with threatened and endangered species. When conducting field work in areas or countries where terminal blood collection is not allowed, the use of restraint devices offer a great alternative for blood collection and can be used with or without anesthesia.

Typically, when anesthesia is used, an anesthetic inhalant gas (commonly Isoflurane) is delivered via nasal cone or induction chamber and maintained at 2-4% concentration using an anesthesia vaporizer and accompanying oxygen tank or oxygen concentrator. Anesthesia may be preferred by some researchers who find it easier to collect the blood sample [13,16], although it has been shown to have no significant difference in survival rates after blood sampling compared to unanesthetized bats [17]. Given that some researchers may choose to use anesthesia, the plunger (Fig. 1b) of this restraint device has been designed to accommodate a ¼” internal diameter vinyl hose, which is commonly used with anesthesia vaporizers. This allows the restraint capsule to additionally function as an induction chamber. If anesthetic gas such as isoflurane vapor is used, additional efforts to scavenge and recover the waste anesthetic gas must be made as these restraints are not hermetic. These methods would include performing anesthesia on a downdraft surgery table, in a chemical fume hood, canopy hood, chemical exhaust snorkel, or other gas scavenging systems.

Second to rodents, bats are the most numerous and diverse group of mammals. Due to this large diversity a blood collection restraint platform that is scalable in size to potentially accommodate all bat species is essential. The base of this restraint is currently designed to accommodate multiple sizes of capsules from a Little Brown bat (*Myotis lucifugus*), up in size to an Egyptian fruit bat (*Rousettus aegyptiacus*). Alternative needs for bat restraints include bat banding for migration and longevity studies, as well as wing punches for genetic analyses.

## Acknowledgements

The authors would like to thank the participants who used the bat restraint devices and provided survey responses, as well as the CDC’s Poxvirus and Rabies Branch and Christina Hutson for supporting the development of prototype bat restraint devices.

## Disclaimer

The findings and conclusions in this report are those of the authors and do not necessarily represent the views of the Centers for Disease Control and Prevention.

